# Differences between the *de novo* proteome and its non-functional precursor can result from neutral constraints on its birth process, not necessarily from natural selection alone

**DOI:** 10.1101/289330

**Authors:** Lou Nielly-Thibault, Christian R Landry

## Abstract

Proteins are among the most important constituents of biological systems. Because all proteins ultimately evolved from previously non-coding DNA, the properties of these non-coding sequences and how they shape the birth of novel proteins are also expected to influence the organization of biological networks. When trying to explain and predict the properties of novel proteins, it is of particular importance to distinguish the contributions of natural selection and other evolutionary forces. Studies in the field typically use non-coding DNA and GC-content-based random-sequence models to generate random expectations for the properties of novel functional proteins. Deviations from these expectations have been interpreted as the result of natural selection. However, interpreting such deviations requires a yet-unattained understanding of the raw material of *de novo* gene birth and its relation to novel functional proteins. We mathematically show how the importance of the “junk” polypeptides that make up this raw material goes beyond their average properties and their filtering by natural selection. We find that the mean of any property among novel functional proteins also depends on its variance among junk polypeptides and its correlation with their rate of evolutionary turnover. In order to exemplify the use of our general theoretical results, we combine them with a simple model that predicts the means and variances of the properties of junk polypeptides from the genomic GC content alone. Under this model, we predict the effect of GC content on the mean length and mean intrinsic disorder of novel functional proteins as a function of evolutionary parameters. We use these predictions to formulate new evolutionary interpretations of published data on the length and intrinsic disorder of novel functional proteins. This work provides a theoretical framework that can serve as a guide for the prediction and interpretation of past and future results in the study of novel proteins and their properties under various evolutionary models. Our results provide the foundation for a better understanding of the properties of cellular networks through the evolutionary origin of their components.

## 1. Introduction

Theoretical and empirical studies of how species acquire new proteins have described several mechanisms with distinct impacts on genomes. Most of these mechanisms, such as gene duplication (Innan and Kondrashov, 2010), horizontal gene transfer (Soucy et al., 2015) and gene fusion (Di Roberto and Peisajovich, 2014), produce novelty by tweaking and rearranging pre-existing gene sequences. However, the phenomenon of *de novo* gene birth departs from this principle, since it consists in the emergence of new genes from sequences that were ancestrally non-genic, or ancestrally non-coding in the case of novel coding genes (McLysaght and Hurst, 2016). Although *de novo* gene birth was once thought to be highly improbable (Jacob, 1977), lineage-specific genes and proteins are observed in a variety of eukaryotes (McLysaght and Guerzoni, 2015), bacteria (Neuhaus et al., 2016) and endosymbiotic organelles (Breton et al., 2011), which suggests that the contribution of *de novo* gene birth to proteomic evolution is not negligible. The biological activities of these novel sequences are often obscured by their lack of homology to other genes, but some of them have been shown to play important and even vital biological roles (Chen et al., 2010; Heinen et al., 2009; Reinhardt et al., 2013). Since *de novo* gene birth is the only source of novel protein families and thus the only way to bring totally novel elements to protein-based cellular networks, it may have significantly influenced the diversity of existing protein structures. For instance, it may be one of the reasons why known proteins seem to form independent homology superfamilies whose estimated times of origination are scattered across the whole history of life (Edwards et al., 2013).

The definition of a functional sequence is important in the study of *de novo* genes, as it distinguishes them from the raw material of *de novo* gene birth, i.e. the “spurious” transcription and translation of non-genic sequences (Wilson and Masel, 2011). Within an organism, a given structure (e.g., a DNA sequence) can have various effects on the phenotype, some of which may be beneficial (i.e. increase organismal fitness). Among these beneficial effects, some may have played a role in the conservation of the structure by natural selection up to the present. That is to say, those effects made the structure much more likely to persist. In evolutionary biology, such beneficial effects that contributed to the conservation of a structure are called the functions of this structure (Doolittle et al., 2014). This is the definition of “function” that we use in this work, and it should not be confused with definitions used in genetics and biochemistry, which are closer to the notion of effect (Kellis et al., 2014). Since the effect of a structure must already have impacted its evolution to qualify as a functional effect, a new function must necessarily have previously existed as a non-functional effect. We use the term “functionalization” to refer to the acquisition of the status of function by a pre-existing effect via its contribution to the conservation of the associated structure. With these definitions, a polypeptide can be seen as an effect of the open reading frame (ORF) that encodes it, and the *de novo* emergence of a polypeptide-coding gene corresponds to the functionalization of a polypeptide. We use the term “polypeptide” to refer to a chain of amino acids of any length, since the terms “peptide” and “protein” carry connotations in this regard. The process of polypeptide functionalization requires the existence of non-functional polypeptides, or junk polypeptides (JPs), since only pre-existing non-functional effects can undergo functionalization. JPs are non-functional because selection for their expression and their structure did not yet contribute to the conservation of their ORFs, but this does not mean that they are not beneficial. A positive impact on fitness is in fact necessary, but not sufficient, for a JP to functionalize, because this positive impact could be too weak to cause the conservation of the JP in the face of genetic drift and mutation (Ohta, 1992). Although the expression of a single JP is presumably unlikely to be strongly beneficial, the “testing” of a large diversity of JPs during evolution increases the chance that some will functionalize.

The conceptual distinction between JPs and functional polypeptides is relevant in evolutionary proteomics, since JPs have not been shaped by positive selection for any activity and are the raw material from which natural selection can draw. As a result, evolutionary models that explain their structural and regulatory properties will likely not apply to functional polypeptides, and vice versa. For similar purposes, functional polypeptides can be meaningfully divided into novel functional polypeptides (novFPs), which recently functionalized and are identical to their non-functional ancestral forms, and ancestral functional polypeptides (ancFPs), which have been altered by evolutionary forces since their functionalization. As most of the canonical coding genes (ORFs annotated by genome databases) have divergent homologs in multiple species, it is safe to assume that they largely meet the definition of ancFPs. The distinction between novFPs and JPs is more difficult to make and remains the most important one to investigate.

This classification of the whole proteome into three groups of polypeptides leads to the division of proteomic evolution into three phases: the turnover of JPs, which is devoid of effective positive selection for polypeptide activities; functionalization, which produces novFPs by filtering JPs without modifying them; and the subsequent modification, loss, duplication and fusion of novFPs and ancFPs (fig. 1). Since well-studied polypeptides are mostly ancFPs, the last two phases have been the objects of most research in evolutionary proteomics. Studying the evolutionary turnover of JPs and their functionalization should thus be complementary to our current understanding of proteome evolution.

**Fig 1.**
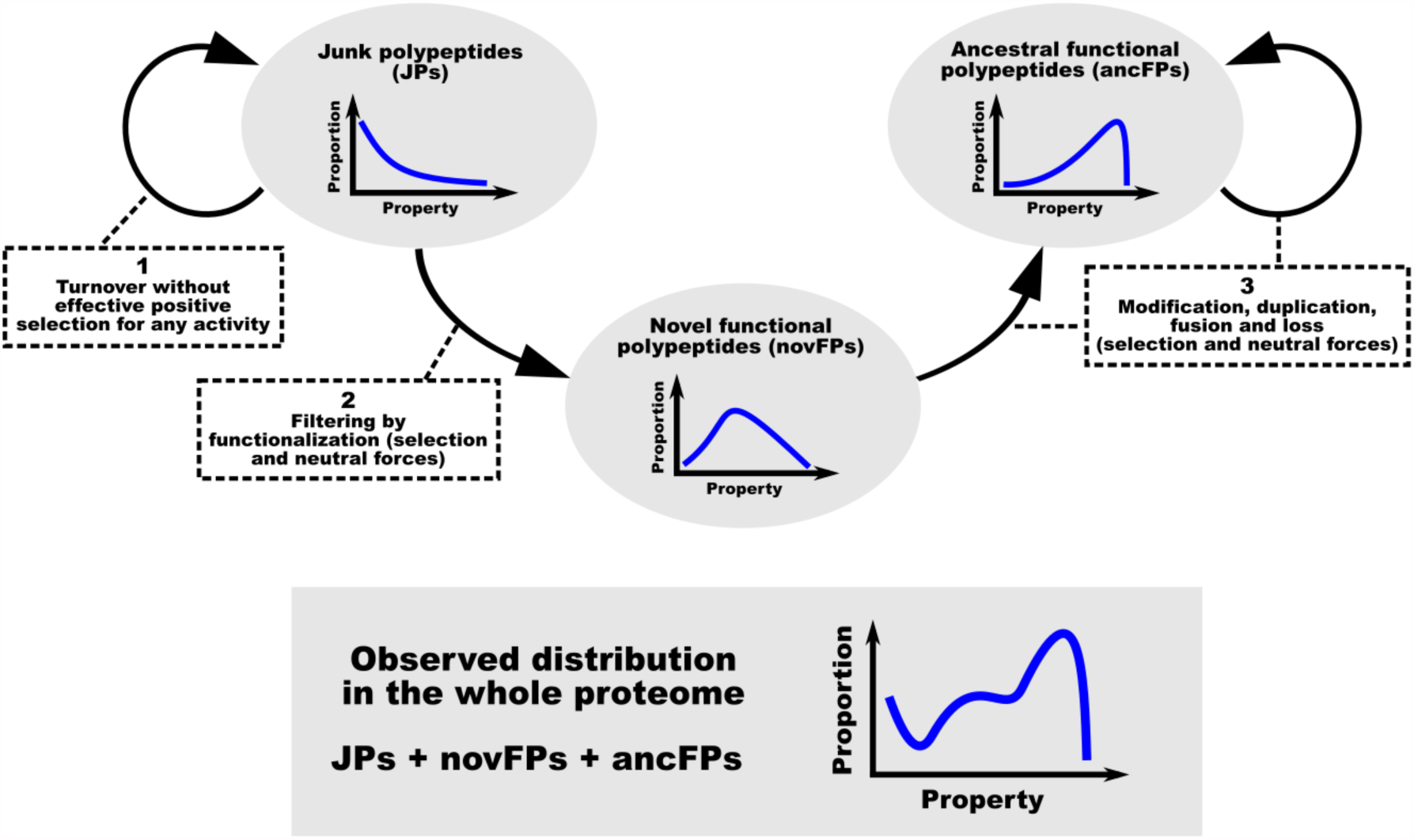
Conceptual classification of polypeptides and their phases of evolution with respect to functionalization. The curves describe the distribution of a hypothetical unidimensional property of polypeptides within each class. A JP is a polypeptide that did not yet contribute to the evolutionary conservation of the ORF that encodes it (i.e. it has not functionalized). A novFP is the immediate product of functionalization: a functional polypeptide that is identical to its ancestral JP. An ancFP is a functional polypeptide that is no longer identical to its ancestral JP. All aspects of the distributions shown are intended as arbitrary examples. For instance, novFPs may not be intermediate between JPs and ancFPs in some regards, and their relative contributions to the proteome-wide distribution may vary.

Although many authors agree that conservation by natural selection should be part of the definition of *de novo* genes (McLysaght and Hurst, 2016; Schlotterer, 2015), the exact moment in the existence of a polypeptide at which *de novo* gene birth happens has not been agreed upon, which makes “*de novo* gene birth” and related terms confusing in practice. For clarity, we hereinafter avoid these terms and instead use the above-defined concepts of JP, novFP, ancFP and functionalization to describe the evolution of proteomes. However, it is worth noting that polypeptides which are called “*de novo*” or “novel” often correspond to novFPs and relatively young ancFPs (McLysaght and Hurst, 2016) but sometimes include JPs (Lu et al., 2017), while the term “protogene” seems to encompass ORFs encoding JPs, novFPs and young ancFPs (Carvunis et al., 2012).

The set of all JPs expressed by a population, which we call the junk proteome, can be seen as a collection of fixed or segregating alleles across a set of loci. There is direct evidence for the existence of this proteome: experimental studies have shown that in a variety of organisms, a large part of intergenic DNA is transcribed into 5’-capped and polyadenylated transcripts (Jensen et al., 2013) which can be translated (Ingolia et al., 2014; Ruiz-Orera et al., 2014). Contrary to canonical genes, these transcripts show signs of suboptimal translation (Guttman et al., 2013) and rapid evolution (Neme and Tautz, 2016). Additionally, the so-called untranslated regions (UTRs) of canonical transcripts and the alternative reading frames within canonical ORFs are sometimes translated into polypeptides that lack known functions (Ingolia et al., 2014; Landry et al., 2015; Mouilleron et al., 2016; Vanderperre et al., 2013) and may thus be JPs. In mice, many translated ORFs in protein-coding genes and long non-coding RNAs were shown to evolve without any detectable selective constraints on the polypeptides that they encode (Ruiz-Orera et al., 2018). Experiments have shown that the fitness effect of the expression of random DNA sequences in bacteria is often slightly beneficial (Neme et al., 2017), although sequences explored in such studies may not reflect those explored by the natural evolution of JPs.

Several studies have inferred lineage-specific functional polypeptides (i.e. novFPs and young ancFPs) and compared them with ancient ancFPs, *in silico* translations of non-coding DNA and randomly generated polypeptides. Assuming that the young ancFPs inferred by those studies are largely similar to novFPs, their results suggest that novFPs typically differ from ancient ancFPs by their short length, weak expression (Schlotterer, 2015), peripheral position in cellular networks and random-sequence-like secondary structure (Abrusán, 2013). It has been proposed that JPs and novFPs may be largely shaped by the genomic GC content through its effects on the properties of ORFs occurring in non-coding DNA sequences (Ángyán et al., 2012). Correlations supporting this role of GC content were observed for many quantities computed from the sequences of ORFs encoding inferred novFPs, although the averages of these quantities often depart from random expectations based on GC content (Basile et al., 2017). Such discrepancies were previously interpreted as the result of natural selection (Wilson et al., 2017), which is in line with the intuition that the probability of functionalization of a beneficial JP increases with its positive effect on fitness. However, several aspects of polypeptide functionalization require clarification before we can confidently interpret the average properties of observed novFPs and their differences from random or non-coding sequences.

In this article, we derive general mathematical results that link the average properties of novFPs to those of JPs. We find that the absolute difference in the mean of a polypeptide property between JPs and novFPs is proportional to its standard deviation among JPs. Furthermore, we show that the average of a property among JPs may not be an appropriate neutral expectation for the corresponding average among novFPs, since a difference between these two values can result from the correlation of the property with the rate of evolutionary turnover of JPs, their selection coefficient, or both. To illustrate how our general equations can be used to study particular polypeptide properties under particular models, we combine them with a GC-content-based random-sequence model of JPs to predict how the genomic GC content and evolutionary parameters interact to determine the mean length and mean intrinsic structural disorder of novFPs.

## 2. RESULTS

### 2.1 The difference in the average of a property between JPs and novFPs is proportional to the standard deviation of this property among JPs

We consider the evolution of the proteome in a single species over an arbitrary time period (e.g., a single branch in a phylogenetic tree). We compare two distributions on the space of possible polypeptides: the average of the junk proteome over this time period (where the relative frequency of each possible category of polypeptides is an average over time) and all novFPs that emerge by functionalization during the same time period. We simply refer to these two distributions as JPs and novFPs, respectively. To compare JPs and novFPs, we use the general concept of quantitative polypeptide property, which we define as a numeric variable that takes a single value in each possible polypeptide-expressing allele. Such properties include, for instance, the length of the polypeptide, the prevalence of some amino acid in its sequence and its expression level. In a group of polypeptides, quantitative polypeptide properties have distributions whose summary statistics can be used to compare JPs and novFPs. Based on the fact that any novFP must first exist as a JP before functionalizing, we find that the difference in the mean of any quantitative polypeptide property *q* between novFPs and JPs is given by the following equation:

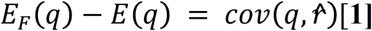

where *E*(*x*) is the expected value (the mean) of a variable, *cov*(*x, y*) is the covariance of two variables, the subscript *F* specifies that a statistic describes novFPs rather than JPs, and 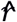 is a quantitative polypeptide property representing the factor by which the relative frequency of each polypeptide changes from JPs to novFPs (see Supplementary Information for details). This equation is analogous to the Robertson-Price identity from quantitative genetics (Lynch and Walsh, 1998; Price, 1970; Robertson, 1966), which states that during a round of natural selection in a population, the mean of a quantitative phenotypic trait changes by an amount equal to the initial covariance of this trait with relative fitness. This analogy between functionalization and natural selection is due to the fact that they both involve the comparison of the relative abundances of types between two populations, where the first population (e.g., JPs or pre-selection individuals) features all the types that are present in the second one (e.g., novFPs or post-selection individuals) and possibly more. This is because both functionalization and natural selection act on pre-existing material but cannot produce novelty on their own. For any process that meets this requirement, the Radon-Nikodým theorem (Yeh, 2006) implies that the difference in the mean of a type-specific quantity (e.g., a quantitative polypeptide property or a quantitative trait) between the second population and the first is equal to the covariance, in the first population, of this quantity with the factor by which the relative abundance of a type changes between the populations (e.g., 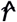 or relative fitness).

To better illustrate the influence of the properties of JPs on those of novFPs, equation 1 can be transformed into:

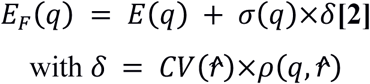

where *σ*(*x*) is the standard deviation of a variable, *CV*(*x*) is its coefficient of variation (the ratio of the standard deviation to the mean) and *ρ*(*x, y*) is the Pearson correlation coefficient of two variables. Because of the mathematical properties of *ρ*, the value of *δ* does not depend on the mean and variance of *q* among JPs, but rather on its relation with functionalization as symbolized by 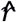.

Equation 2 has implications for the use of random and non-coding controls in the study of novFPs. Such controls were often used to compute expected means and other measures of central tendency for polypeptide properties (Abrusán, 2013; Ángyán et al., 2012; Basile et al., 2017; Wilson et al., 2017). According to equation 2, the standard deviations of the properties of control sequences could be just as useful as their means for predicting the properties of novFPs, provided that the control is representative of real JPs. When comparing the mean of a polypeptide property in such a representative control with the corresponding mean among inferred novFPs, these two means can be used in equation 2 along with the standard deviation in the control to estimate the *δ* of this property. *δ* captures the strength of biases of *de novo* gene birth in favour of certain polypeptide properties without being defined by their distribution among JPs. Thus, in order to interpret average differences between JPs and novFPs given the distributions of the properties of JPs, we need to decompose *δ* into contributions from different evolutionary forces.

### 2.2 Neutral evolutionary forces can cause discrepancies between JPs and novFPs through the rate of evolutionary turnover of JPs

We sought to transform the definition of *δ* from equation 2 into readily interpretable equations. Once we make the additional assumption that JPs are at evolutionary equilibrium, i.e. each JP is gained by mutation as often as it exits the junk proteome by loss or functionalization, then *δ* becomes:

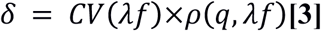

where *λ* is the polypeptide-specific rate of evolutionary turnover (the inverse of the expected time from the appearance of a specific JP by mutation to either its loss or its functionalization) and *f* is the polypeptide-specific probability that a new JP will functionalize before it disappears from the population. Because of the mathematical properties of *CV* and *ρ, δ* is insensitive to the scales of *q, λ* and *f*. As a result, each of them can be replaced with a directly proportional quantity without changing the value of *δ*, which may help to model this value and to estimate it from the observed properties of JPs. For instance, if a model of the evolution of JPs assumed that the turnover rate of a JP is directly proportional to the GC content of its ORF, *λ* could be simply replaced by this variable in equation 3.

While equation 3 could be used to compute *δ* given enough data or assumptions about the junk proteome and its evolution, it does not highlight intuitive possible explanations for the discrepancies between novFPs and JPs, i.e. cases where *δ* is not zero. In order to do this, and without adding any assumption to those behind equation 3, we obtained the following expression of *δ* using equivalence properties of covariance.

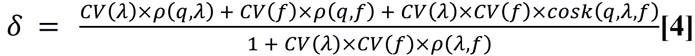

where 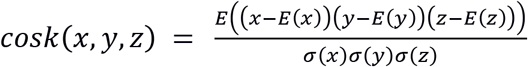 is the coskewness of three variables.

The second term of the numerator in equation 4 confirms previous intuitions about the probability of functionalization: all else being equal, an increase in its correlation with a given polypeptide property results in an increase of this property’s *δ* and thus of its mean among novFPs. More surprisingly, the first term of the numerator indicates that the same relation exists between *δ* and the rate of evolutionary turnover. This entails that the mean of a polypeptide property that has no selective effect on the probability of functionalization can still be different between JPs and novFPs if this property correlates with the rate of evolutionary turnover of JPs. For example, if the turnover of long JPs is especially fast because they mutate at a frequency proportional to their length, this could contribute to their over-representation among novFPs relative to the junk proteome, even without their selection coefficients being larger than those of shorter JPs. In other words, the frequency of successes (events of functionalization) depends as much on the frequency of trials (the turnover rate) as on the probability of success for a single trial (the probability of functionalization). As a result, observed differences between the average properties of novFPs and those of random sequences should be either shown or explicitly assumed not to be caused by neutral biases in turnover rate before they are interpreted as the results of natural selection.

The coskewness that appears in the third term of the numerator in equation 4 is a measure of how any of three variables linearly affects the linear relation between the two others (see Supplemental Information). Like *CV* and *ρ*, it is insensitive to the replacement of a variable by a directly proportional quantity. Despite the diffuculty of its interpretation, it could be estimated from data on the turnover rate and functionalization probability of JPs, or predicted from a model of their evolution.

The denominator of equation 4 is strictly positive and indicates that the overall correlation between the turnover rate and the probability of functionalization negatively affects the magnitude of *δ*. Interestingly, this term does not involve *q*, which means that its value is the same for every polypeptide property in a given species. It can be thought of as a measure of the overall tendency of the junk proteome to preferentially explore polypeptides that are likely to functionalize. It constitutes a baseline to which each source of evolutionary bias represented in the numerator must compare favourably in order to have a strong effect on the average properties of novFPs.

Length and secondary structure are biologically relevant aspects of polypeptides that can be studied *in silico* in arbitrary sequences, which makes them ideal targets for the modelling of JPs and novFPs. In particular, the ISD of novel polypeptides has been a recurrent topic in previous studies (Ángyán et al., 2012; Basile et al., 2017; Wilson et al., 2017). To exemplify the usefulness of our general results, we used them to model the mean length and mean ISD of novFPs given a simple model of the sequences of JPs. This model assumes that all the nucleotides that encode the junk proteome have independently evolved to equilibrium under uniform and strand-symmetric evolutionary pressures, and that a random subset of the ORFs appearing in the resulting sequences are translated into JPs. By combining these assumptions, the mean and standard deviation of any sequence property among JPs can be predicted from a single parameter: the GC content of DNA.

We predicted the means and standard deviations of length and ISD among JPs as functions of the GC content. In the case of length, these predictions were made analytically, while in the case of ISD, we used the model to simulate 100 000 polypeptide sequences for each GC content in a large range of values and we estimated their individual ISD levels with the sequence-wide average of IUPred “long” disorder (Dosztányi et al., 2005). We then applied equation 2 to the resulting means and standard deviations to compute the expected means of length and ISD among novFPs as functions of *δ* and the GC content (fig. 2).

**Fig 2.**
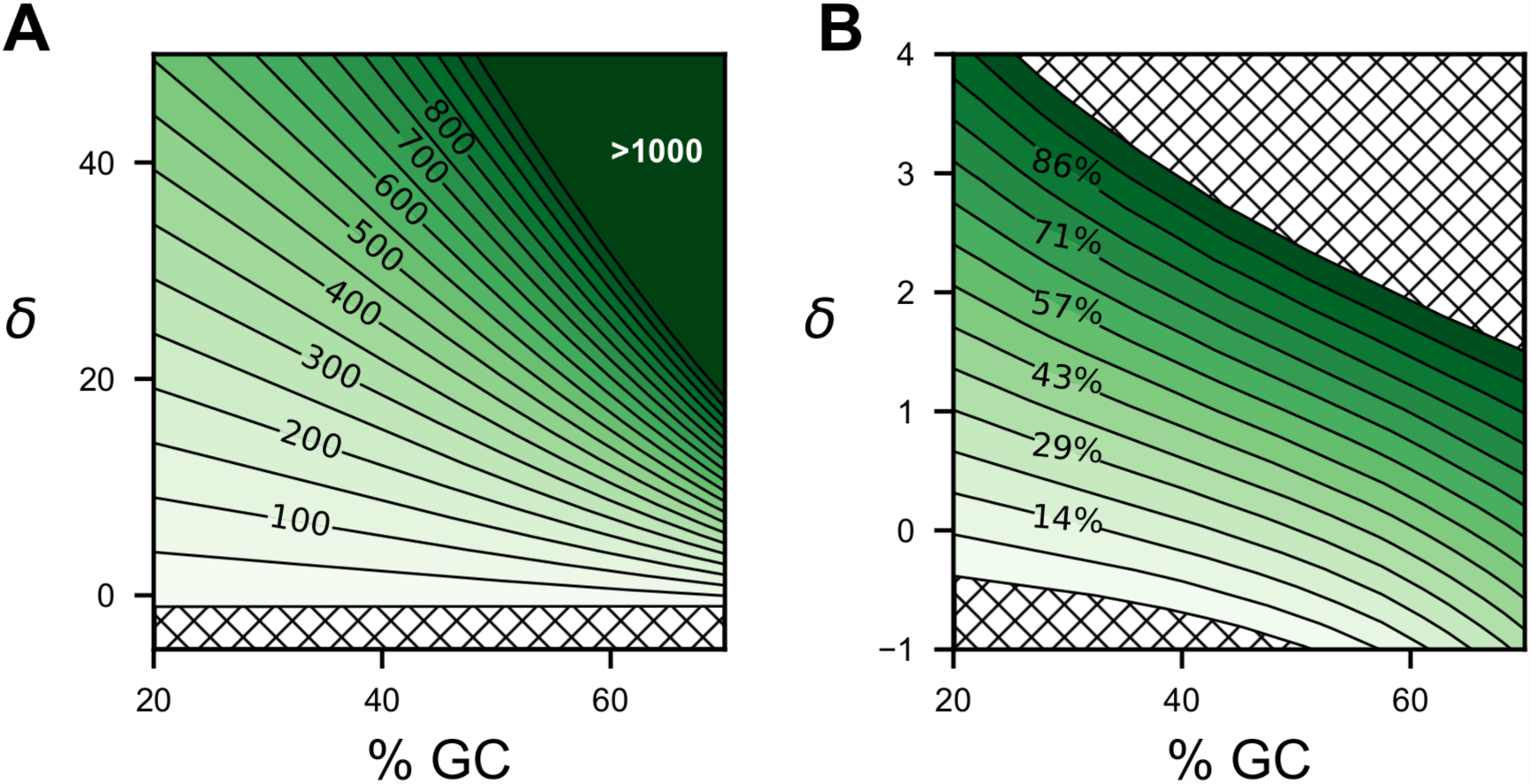
Contour plots of predicted means for the length and intrinsic structural disorder of novel functional polypeptides as functions of *δ* and the GC content. (*A*) Polypeptide length in amino acid residues. (*B*) Sequence-wide mean of the IUPred “long” prediction of intrinsic structural disorder (Dosztányi et al., 2005). Intensity values associated with contour lines are equidistant. As in any contour plot, the vertical distance between contour lines is inversely proportional to the vertical rate of change in intensity. In these specific contour plots, this vertical rate is the standard deviation of the considered property among JPs (equation 2). Since this standard deviation is constant for a given GC content, the contour lines are vertically equidistant. Hatched areas indicate impossible scenarios, that is, negative polypeptide lengths and percentages outside the 0%-100% interval.

In both landscapes of figure 2, the curve obtained by taking an horizontal “slice” at *δ* = 0 corresponds to the relation between the mean properties of JPs and the GC content under the GC-content-based random-sequence model. These horizontal slices at *δ* = 0 are consistent with known effects of an increase in GC content on random polypeptides, namely an increase in their length and their ISD (Basile et al., 2017), even though this is not clearly visible in (fig. 2A). However, figure 2 also shows that for a polypeptide property whose *δ* is non-zero, the effect of GC content on novFPs (the change in tone associated with a horizontal shift) can differ from its effect on random sequences (the result of the same shift at *δ* = 0), since contour lines are curved. It is theoretically possible for these effects to differ in sign as well as magnitude (the same contour line could be decreasing for some GC contents and increasing for others), although this is not the case for the two properties presented here. As implied by equation 2, such inconsistencies of the effect of GC content between JPs and novFPs would be due to the fact that the GC content affects both the standard deviations and the means of the properties of JPs, possibly in opposite directions. In contour plots such as those of figure 2, the standard deviation among JPs corresponds to the vertical slope of the landscapes and is thus inversely proportional to the vertical distance between contour lines. For example, in the case of polypeptide length (fig. 2A), the tightening of contour lines from left to right indicates that the standard deviation of the length of JPs increases with GC content, making the mean length of novFPs especially sensitive to the associated *δ* in GC-rich genomes.

Even though inferred novFPs tend to be shorter than ancFPs, their average length is usually at least 100 residues (Basile et al., 2017; Neme and Tautz, 2013), which is larger than the expected mean length of JPs for common GC contents (fig. 2A, *δ* = 0). Thus, if polypeptides that were detected and classified as novel are representative of novFPs, the value of *δ* associated with polypeptide length is likely to be greater than zero in many species. As explained in our interpretation of equation 4, this would not necessarily mean that a long polypeptide is typically more likely to functionalize than a shorter one. The fact that ancFPs tend to be larger than novFPs is also not conclusive evidence for such a selective advantage of length among JPs, since the evolution of ancFPs may be channelled towards long polypeptides that are very different from JPs of the same length. One intuitive alternative explanation for novFPs being longer than JPs is that since the rate of neutral loss of an ORF is proportional to its length, the length of JPs is positively correlated with their rate of turnover, which increases the *δ* of polypeptide length as shown in equation 4. However, the strength of this correlation depends on how much variation in the rate of turnover is caused by factors other than ORF length (such as the turnover of promoter sequences), and its contribution to *δ* also depends on the overall correlation between the rate of turnover and the probability of functionalization, as shown by the denominator of equation 4. It is therefore currently hard to tell if this effect is strong enough to fully explain the observed shift in mean length between random ORFs and those expressing novFPs, although this point could be clarified by modelling or estimating the turnover rate and probability of functionalization of JPs. Despite the ambiguity as to the causes of the apparent length difference between JPs and novFPs, this difference should increase with the standard deviation of the length of JPs, and thus with GC content, unless this effect cancels out with a decrease of *δ* in GC-rich genomes (fig. 2A). For instance, the mutation spectrum of an organism affects both its genomic GC content and the relation between the sequence of an ORF and its rate of mutation, which could lead to an inter-specific correlation between GC content and *δ* for various polypeptide properties. Implications of the length of JPs for the properties of novFPs have been largely ignored since studies of *de novo* gene birth usually use random or non-coding controls that are intentionally biased against short ORFs (Abrusán, 2013; Ángyán et al., 2012; Basile et al., 2017; Lu et al., 2017; Neme et al., 2017; Wilson et al., 2017). Such practices may be partly justifiable if very short JPs turn out to contribute negligibly to *de novo* gene birth because of slow turnover or low probability of functionalization, but this remains to be shown.

In the house mouse, *in silico* predictions suggest that novFPs have higher ISD than potential polypeptides encoded by intergenic DNA, which was interpreted as a result of natural selection in favour of high ISD during *de novo* gene birth (Wilson et al., 2017). Other results suggest that this trend may be specific to certain organisms and certain values of genomic GC content **(30)**. As the average GC content of house mouse DNA is 42% (Elhaik and Graur, 2014) and the average IUPred long disorder of its novFPs is close to 55% (Wilson et al., 2017), our model predicts that the *δ* of this specific measure of ISD should be above 1 in house mouse (fig. 2B), more precisely 1.23. This value being larger than zero is consistent with the conclusion of (Wilson et al., 2017) that novFPs appear more disordered than the raw material of *de novo* gene birth, assuming that the non-coding control sequences they used are well summarized by a single GC content that is close to 42%. By a similar reasoning, given the 38% GC content observed in yeast DNA (Engel et al., 2014) and the 32% average IUPred long disorder of yeast novFPs reported by (Wilson et al., 2017), the associated value of *δ* should be between 0 and 1 (fig. 2B), more precisely 0.47. Under our GC-content-based model, this suggests that given the GC contents of mouse and yeast genomes, the biases of turnover and functionalization in favour of disordered polypeptides are stronger in mouse than in yeast.

Our positive estimates of *δ* for the sequence-wide average of IUPred long disorder in mouse and yeast may reflect positive correlations of this polypeptide property with the turnover rate and/or the functionalization probability (see equation 4). However, IUPred “long” disorder is an estimator of ISD and we can only assume that the landscape of the actual average proportion of disordered amino acid residues in novFPs is similar to (fig. 2B) under the GC-content-based model. As a warning against this assumption, the corresponding landscape computed from IUPred “short” disorder (supplementary fig. S1) is different from (fig. 2B) in terms of the magnitude of *δ* because the means and standard deviations of “long” and “short” predictors of ISD among JPs have different relations to the GC content, even though they are both meant to estimate the proportion of disordered residues. Nevertheless, the difference between mouse and yeast in the estimated *δ* for the same measure of ISD suggests that the difference in average ISD between their novFPs is not solely driven by the mean and standard deviation of ISD among JPs, but also by a difference in the relation of ISD with the turnover rate and the probability of functionalization of JPs. Future studies may reveal that some components of *δ* can be predicted from the GC content, which would make the latter even more useful than figure 2 suggests for the prediction of inter-specific differences in the average properties of novFPs.

## 3. Discussion and conclusions

When making inferences such as those presented in figure 2, the use of equation 2 is inherently valid, since this equation stems from the definitions of JPs and novFPs. However, the means and standard deviations of the properties of JPs, which are needed to apply equation 2, are model-dependent. Therefore, the type of predictions that we made as to the value of *δ* under a GC-content-based model may not apply to organisms where JPs are not well described by such a model. For instance, mammalian genomes are known to be organized into compositional domains with various GC contents (Elhaik and Graur, 2014). In a study in yeast, candidate *de novo* genes had a significant tendency to be located in GC-riched regions of the genome (Vakirlis et al., 2017). If several different GC contents contribute to a single junk proteome, the means and standard deviations of the properties of JPs may be different from those expected under our random-sequence model given the average GC content. It is however possible that such a model will apply to each compositional domain separately, in which case the junk proteome would be readily modelled by drawing values of GC content from an appropriate distribution and using each value to generate a random polypeptide. From there, equations 1, 2, 3 and 4 would apply just as they did for the simpler case of a single GC content.

Although the predictions that we made using a random-sequence model of JPs only involve their sequence and structure, intrinsic aspects of their expression may also be understood as quantitative polypeptide properties and analyzed using equations presented here. Transcription and translation levels of JP-encoding ORFs seem especially relevant since, as they approach zero, the probability that a JP functionalizes also goes towards zero and its other properties become irrelevant. Since transcription and translation are controlled by local sequence elements, knowledge of these elements may eventually be combined with a random-sequence model to predict the regulatory properties of JPs, like we did for their sequence properties. Studies of the transcriptional activity of synthetic random DNA in *Escherichia coli* (Yona et al., 2017) and yeast (Boer et al., 2018) show that such sequences frequently contain the patterns required for the initiation and regulation of transcription. Factors that are external to intergenic regions also seem to play a role in the expression of JPs and their functionalization, such as bidirectional promoters (Vakirlis et al., 2017), translated UTRs and translated alternative ORFs within canonical ORFs (Vanderperre et al., 2013). Although understanding the importance of these factors may require more than a simple random-sequence model, their impacts on JPs will be “inherited” by novFPs in accordance with the general equations that we developed.

The determinants of the properties of polypeptides resulting from *de novo* gene birth were previously studied empirically by comparing them to random and non-coding sequences (Abrusán, 2013; Ángyán et al., 2012; Basile et al., 2017; Lu et al., 2017; Wilson et al., 2017), but the field lacked the theoretical tools to interpret observations in terms of evolutionary forces. We have defined a classification of polypeptides and their evolutionary history (fig. 1) that clarifies the process of *de novo* gene birth sufficiently to link the properties of its raw material to those of its products through broadly applicable equations. These equations show how the mean of a quantitative polypeptide property among the products of *de novo* gene birth depends on its mean, its standard deviation and its relation to both the probability of functionalization and the turnover rate, which suggests potential roles for both natural selection and neutral forces. We also showed how a simple GC-content-based model of non-functional polypeptides can be combined with our general theoretical framework to infer evolutionary parameters of *de novo* gene birth from the properties of its products, or vice versa. Although our results specify how knowledge of the structure, expression and evolution of the non-functional proteome can be used to explain and predict the properties of novel functional polypeptides, much of this knowledge remains to be uncovered by further empirical, experimental and theoretical investigation.

## 4. MATERIALS AND METHODS

### 4.1 A general model of the link between the properties of JPs and those of novFPs

The details of all formal reasonings can be found in Supplemental Information. We defined an average of the junk proteome over time (a probability measure on the space of possible polypeptides), such that any category of polypeptides that has a frequency of zero in this average statistical population also has a frequency of zero among novFPs. We then used established theorems from measure theory and probability theory to compare the mean of an arbitrary polypeptide property between JPs and novFPs, which led to equations 1 and 2. We then made the additional assumption the JPs are at evolutionary equilibrium, and thus the rate of appearance of any JP by mutation is equal to its rate of loss. This assumption implies that the intrinsic rate of loss of JPs from a given category can be used as an indicator of the rate at which evolution explores this category. By combining the assumption of evolutionary equilibrium with our definition of *δ* from equation 2, we obtained equations 3 and 4.

### 4.2 A random-sequence model of the properties of JPs

Mathematical developments and results that parallel this section are presented in Supplemental Information. To quantitatively model the properties of JPs, we made five assumptions about the DNA encoding them: 1) all sites evolve independently, 2) the transition probability matrix is constant across sites, 3) the transition probability matrix is the same on both strands, 4) each site has reached evolutionary equilibrium, and 5) a random subset of ATG codons define ORFs that are translated into JPs. Assumptions 1 and 2 allow us to focus on a single site and generalize our findings to the whole sequence. Assumptions 3 and 4 entail that if two nucleotides are Watson-Crick complements, then a given site is equally likely to display either of them. As a result, complementary nucleotides are equally frequent within and between strands, and the frequency of each of the four nucleotides is a function of GC content. Since sites are independent, GC content is the only parameter needed to predict probability distributions for the properties of randomly occurring ORFs under this model. Assumption 5 allows us to extend our predictions to the properties of JPs expressed from those ORFs.

Predicting the length distribution of JPs is equivalent to predicting the distribution of the number of in-frame sense codons separating each ATG codon from the closest downstream in-frame stop codon. Since, in our model, consecutive non-overlapping DNA 3-mers are statistically independent and have the same probability of being stop codons, the number of sense codons in an ORF follows a geometric distribution. The only parameter of this distribution is the frequency of stop codons, which is a function of GC content under our model. We thus predicted the exact shape of the length distribution of JPs as a function of GC content.

### 4.3 Predicting the mean length and mean ISD of novFPs from the GC content and *δ*

We combined equation 2 with the properties of geometric distributions to compute the landscape of the average length of novFPs as a function of GC content and *δ* (fig. 2A). In order to obtain the landscapes of average IUPred long (fig. 2B) and short (supplementary fig. S1) disorders among novFPs, we randomly generated the sequences of 100 000 JPs for each value of GC content from 20% to 80% with steps of 2.5%. We then computed the per-amino-acid “long” and “short” disorder scores using IUPred (Dosztányi et al., 2005), averaged the two types of scores separately within each sequence, computed the mean and standard deviation of the sequence-wide average of each score for each GC content, and applied equation 2 to those means and standard deviations to compute the landscapes.

## ACKNOWLEDGMENTS

L.N.T. is supported by an Alexander Graham Bell PhD fellowship from the Natural Sciences and Engineering Research Council (NSERC). C.R.L. is supported by a Discovery grant from the NSERC and a Team grant from the Fonds de recherche du Québec – Nature et technologie (FRQNT) and holds the Canada Research Chair in Evolutionary Cell and Systems Biology. The authors thank C.K. Griswold and H. Martin for their comments on the manuscript.

---

**Supplementary fig. S1.**
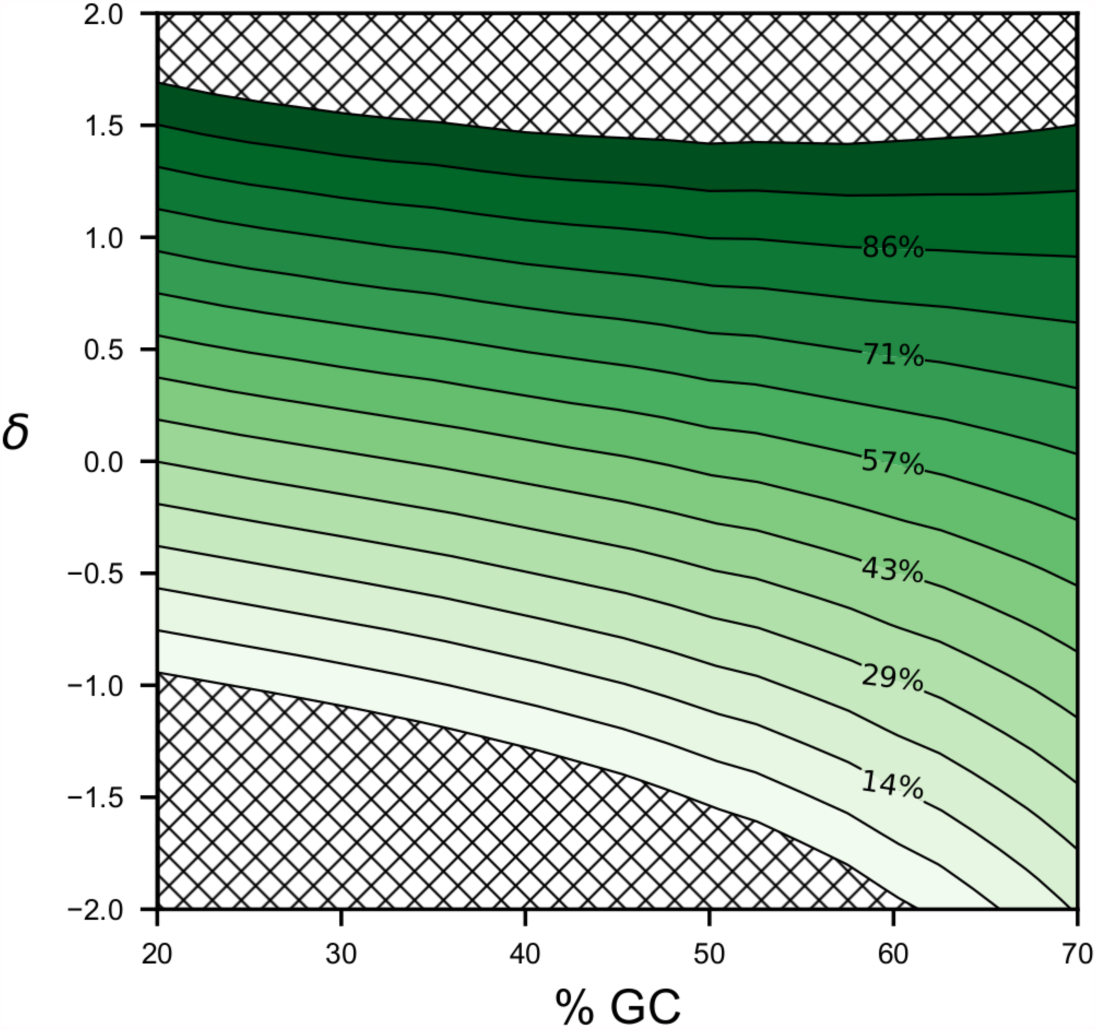
Contour plot of the predicted mean IUPred short disorder of novel functional polypeptides as a function of *δ* and the GC content. Intensity values associated with contour lines are equidistant. As in any contour plot, the vertical distance between contour lines is inversely proportional to the vertical slope of the landscape. In this contour plots, the vertical slope is the standard deviation of the considered property among JPs (equation 2). Since this standard deviation is constant for a given GC content, the contour lines are vertically equidistant. Hatched areas indicate impossible percentages, i.e. outside the 0%-100% interval.

## SUPPLEMENTARY METHODS

### 1. A general model of the link between the properties of JPs and those of novFPs

This section is a detailed reasoning leading to equations 1, 2, 3 and 4. Let *Ω*be the space of all possible polypeptides (or polypeptide-expressing alleles). Over a given period of time, the average proportions of the junk proteome belonging to each possible category of polypeptides form a probability measure *P* on *Ω*. In other words, for each subset *S* of *Ω*, *P*(*S*) is the time-averaged proportion of JPs that fall in the polypeptide category *S*, which implies that *P*(*Ω*) = 1. Similarly, the polypeptides that functionalize in the same period of time form a probability measure *P*_*F*_ on *Ω*. For any quantitative polypeptide property *q*, i.e. any function that assigns a number to each possible polypeptide in *Ω*, statistics like the mean and variance of *q*are defined separately for each probability measure. Hereinafter, we use the subscript *F* to distinguish between statistics defined for *P* and those defined for *P*_*F*_. For example, the mean (expected value) of a property *q* among JPs will be denoted by *E*(*q*), while its mean among novFPs will be denoted by *E*_*F*_(*q*).

Because JPs are not modified by their functionalization, the probability measures *P* and *P*_*F*_ have a special relationship: for any subset *S* of *Ω* such that *P*(*S*) = 0, it is also true that *P*_*F*_(*S*) = 0. In other words, any category of polypeptides that occurs in novFPs necessarily occurred in the junk proteome at some point. Because of this relationship between the two measures (*P*_*F*_ is “absolutely continuous” with respect to *P*), the Radon-Nikodým theorem (Yeh, 2006) implies that there exists a quantitative polypeptide property 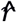 such that, for any subset *S* of *Ω* with *P*(*S*) ≠ 0, the average of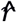among JPs that belong to *S* is given by 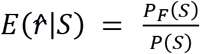. The property 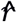 is thus the factor by which the relative frequency of a polypeptide changes from JPs to novFPs, and 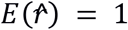. If we define *R* as the total rate of functionalization events over time, *L* as the total number of loci expressing JPs and *r* as the polypeptide-specific ratio of the rate of functionalization events to the time-averaged number of loci expressing this polypeptide, then we have 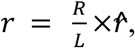, 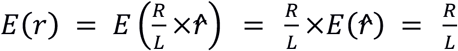 and thus 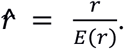.

Given a quantitative polypeptide property, representing its mean among novFPs as a function of its mean among JPs would be useful in the study of *de novo* gene birth. Because of the way we defined 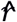 from *P* and *P*_*F*_ (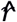 is the Radon-Nikodým derivative of *P*_*F*_ with respect to *P*), it follows (Yeh, 2006) that for any quantitative polypeptide property *q*, we have:

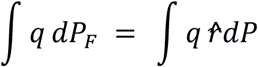

where *∫ × dµ* is the Lebesgue integral of the function *x* with respect to the measure µ.

The expected value of a random variable (a function) is defined as its Lebesgue integral with respect to a probability measure such as *P* or *P*_*F*_. Therefore, the above equation is equivalent to:

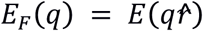

By applying the property of covariance *E*(*xy*) = *E*(*x*)*E*(*y*) + *cov x, y*, we obtain:

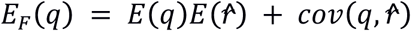

Since 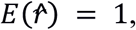, we obtain equation 1:

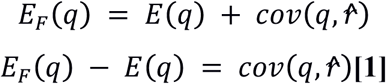

By applying the definition of the Pearson correlation coefficient 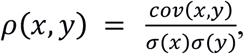, the fact that 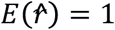 and the definition of the coefficient of variation 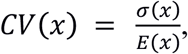, the role of 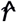 in equation 1 can be concentrated in a single parameter*δ*that is insensitive to the multiplication of 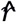 by any positive constant:

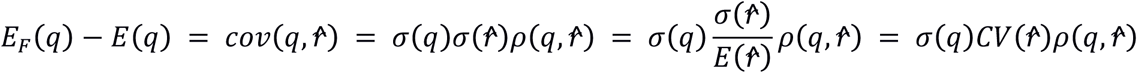

For any *k* > 0:

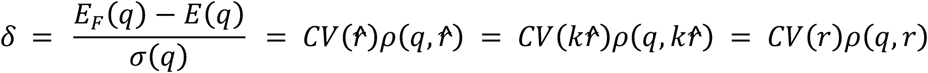

We thus obtain equation 2:

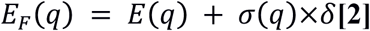

Since the quantitative polypeptide property *r* is the polypeptide-specific ratio of the rate of functionalization of a polypeptide to the average number of non-functional instances of this polypeptide expressed in the species, it can be understood as the product*r* = *γf*, where *γ* is the ratio of the frequency of allele gain (appearance by mutation) to the average number of non-functional loci expressing the polypeptide, and *f* is the probability that such a gain leads to the functionalization of the polypeptide (the probability of functionalization). If we make the additional assumption that the junk proteome is at evolutionary equilibrium, i.e. JPs from any category are gained as often as they are lost or functionalized, then the rate of appearance of each JP is equal to its rate of loss/functionalization. We thus have *r* = *λf*, where *λ* is the ratio of the combined frequency of functionalization and complete loss of a polypeptide (an allele) to the average number of non-functional loci expressing this polypeptide. *λ* can also be interpreted as the rate at which a single JP exits the junk proteome by loss or functionalization (the inverse of its expected lifetime as a JP). Then, from the definition of *δ*, we obtain equation 3:

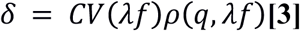

By the definitions of the Pearson correlation coefficient and the coefficient of variation:

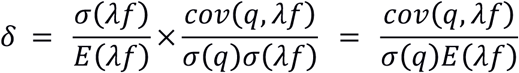

By transforming *E*(*λf*) using the properties of covariance:

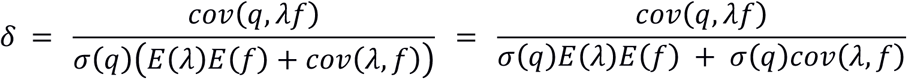

By decomposing the numerator as the covariance of a product of random variables according to **(Bohrnstedt and Goldberger, 1969)**:

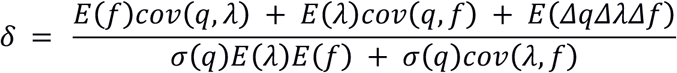

where *Δx* = *x* − *E*(*x*). By dividing the numerator and the denominator with *σ*(*q*)*E*(*λ*)*E*(*f*):

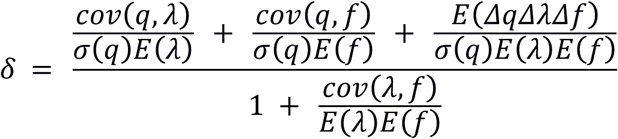

By taking the factor *σ*(*λ*)*σ*(*f*) out of the rightmost term of the numerator:

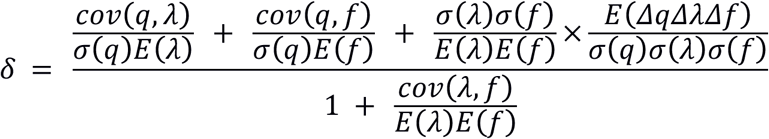

By applying the definition of the Pearson correlation coefficient three times:

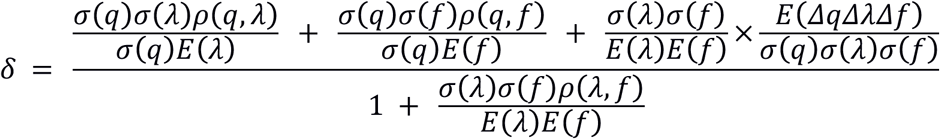

By cancelling and rearranging factors within terms:

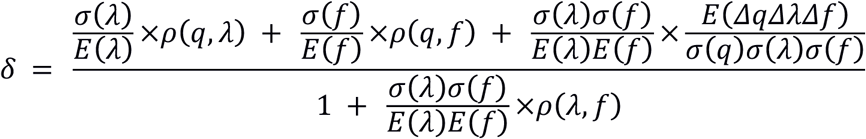

By applying the definition of the coefficient of variation six times:

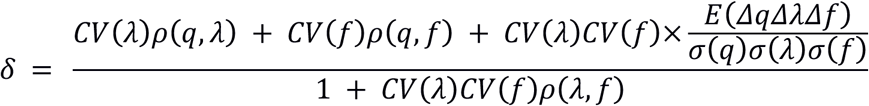

By the definition of the coskewness of three variables 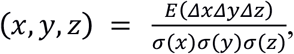, we obtain equation 4:

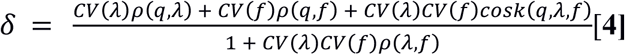

To facilitate the interpretation of coskewness, consider the standard score 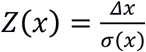 which as a mean of 0 and a variance of 1.

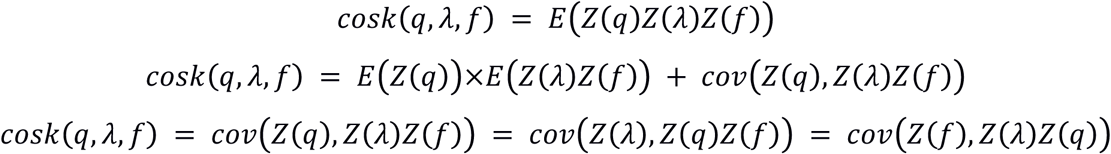

Also consider the fact that *E*(*Z*(*x*)*Z*(*y*)) = *cov*(*Z*(*x*),(*Z*(*y*)) = *ρ*(*x, y*).

While the Pearson correlation coefficient is the mean of the product of the standard scores of two variables, coskewness is the covariance of this same product with the standard score of a third variable. Roughly speaking, coskewness is a measure of how any of the three variables linearly affect the linear relation between the two others.

The denominator in equation 4 is strictly positive, since, by the definition of the coefficient of variation and the properties of covariance:

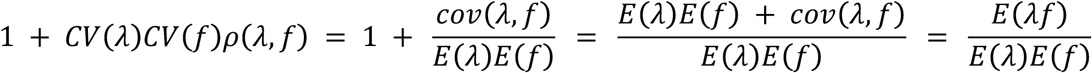

and both *E*(*λf*) and *E*(*λ*)*E*(*f*) are positive.

### 2. Modelling the length distribution of JPs under a GC-content-based random-sequence model

Under our random-sequence model, the frequency *p*_*N*_ of each nucleotide *N* is a function of the GC content, which we denote by *p*_*G/C*_ = *p*_*G*_ + *p*_*C*_. The frequencies of the four nucleotides are given by:

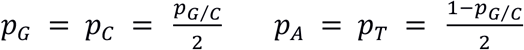

Since, in this model, consecutive non-overlapping DNA 3-mers are statistically independent and have the same probability of being stop codons, the number of sense codons in an ORF follows a geometric distribution with the following probability mass function:

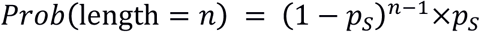

where *n* is any positive integer and *p*_*s*_ is the probability that a DNA 3-mer is a stop codon. Under our assumptions, the frequency of a DNA word is equal to the product of the frequencies of the nucleotides composing it. Using this principle to calculate *p*_*s*_, we get:

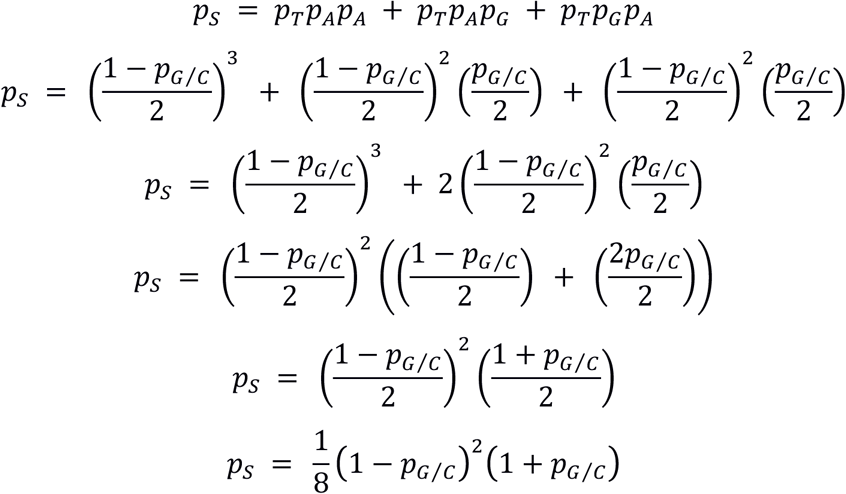

Using the properties of geometric distributions, we obtained the mean and standard deviation of the length of JPs as functions of the frequency of stop codons, which is itself determined by the GC content:

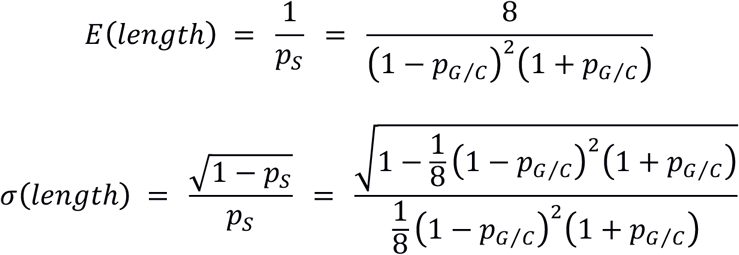

Using these equations in combination with equation 2, we computed the landscape of the mean length of novFPs as a function of the GC content and *δ*, which is shown in figure 2A.

## REFERENCES

Abrusán, G. (2013). Integration of New Genes into Cellular Networks, and Their Structural Maturation. Genetics 195, 1407–1417.

Ángyán, A.F., Perczel, A., and Gáspári, Z. (2012). Estimating intrinsic structural preferences of de novo emerging random-sequence proteins: is aggregation the main bottleneck? FEBS Lett 586, 2468–2472.

Basile, W., Sachenkova, O., Light, S., and Elofsson, A. (2017). High GC content causes orphan proteins to be intrinsically disordered. PLoS Comput Biol 13, e1005375.

Boer, C. de, Sadeh, R., Friedman, N., and Regev, A. (2018). Deciphering cis-regulatory logic with 100 million random promoters. BioRxiv 224907.

Breton, S., Stewart, D.T., Shepardson, S., Trdan, R.J., Bogan, A.E., Chapman, E.G., Ruminas, A.J., Piontkivska, H., and Hoeh, W.R. (2011). Novel Protein Genes in Animal mtDNA: A New Sex Determination System in Freshwater Mussels (Bivalvia: Unionoida)? Mol. Biol. Evol. 28, 1645–1659.

Carvunis, A.R., Rolland, T., Wapinski, I., Calderwood, M.A., Yildirim, M.A., Simonis, N., Charloteaux, B., Hidalgo, C.A., Barbette, J., Santhanam, B., et al. (2012). Proto-genes and de novo gene birth. Nature 487, 370–374.

Chen, S., Zhang, Y.E., and Long, M. (2010). New genes in Drosophila quickly become essential. Science 330, 1682–1685.

Di Roberto, R.B., and Peisajovich, S.G. (2014). The role of domain shuffling in the evolution of signaling networks. J Exp Zool B Mol Dev Evol 322, 65–72.

Doolittle, W.F., Brunet, T.D., Linquist, S., and Gregory, T.R. (2014). Distinguishing between “function” and “effect” in genome biology. Genome Biol Evo. 6, 1234–1237.

Dosztányi, Z., Csizmok, V., Tompa, P., and Simon, I. (2005). IUPred: web server for the prediction of intrinsically unstructured regions of proteins based on estimated energy content. Bioinformatics 21, 3433–3434.

Edwards, H., Abeln, S., and Deane, C.M. (2013). Exploring Fold Space Preferences of New-born and Ancient Protein Superfamilies. Plos Comput. Biol. 9, 11.

Elhaik, E., and Graur, D. (2014). A comparative study and a phylogenetic exploration of the compositional architectures of mammalian nuclear genomes. PLoS Comput Biol 10, e1003925.

Engel, S.R., Dietrich, F.S., Fisk, D.G., Binkley, G., Balakrishnan, R., Costanzo, M.C., Dwight, S.S., Hitz, B.C., Karra, K., Nash, R.S., et al. (2014). The reference genome sequence of Saccharomyces cerevisiae: then and now. G3 Bethesda 4, 389–398.

Guttman, M., Russell, P., Ingolia, N.T., Weissman, J.S., and Lander, E.S. (2013). Ribosome profiling provides evidence that large noncoding RNAs do not encode proteins. Cell 154, 240–251.

Heinen, T.J., Staubach, F., Häming, D., and Tautz, D. (2009). Emergence of a new gene from an intergenic region. Curr Biol 19, 1527–1531.

Ingolia, N.T., Brar, G.A., Stern-Ginossar, N., Harris, M.S., Talhouarne, G.J., Jackson, S.E., Wills, M.R., and Weissman, J.S. (2014). Ribosome profiling reveals pervasive translation outside of annotated protein-coding genes. Cell Rep 8, 1365–1379.

Innan, H., and Kondrashov, F. (2010). The evolution of gene duplications: classifying and distinguishing between models. Nat Rev Genet 11, 97–108.

Jacob, F. (1977). Evolution and tinkering. Science 196, 1161–1166.

Jensen, T.H., Jacquier, A., and Libri, D. (2013). Dealing with pervasive transcription. Mol Cell 52, 473–484.

Kellis, M., Wold, B., Snyder, M.P., Bernstein, B.E., Kundaje, A., Marinov, G.K., Ward, L.D., Birney, E., Crawford, G.E., Dekker, J., et al. (2014). Defining functional DNA elements in the human genome. Proc Natl Acad Sci U A 111, 6131–6138.

Landry, C.R., Zhong, X.F., Nielly-Thibault, L., and Roucou, X. (2015). Found in translation: functions and evolution of a recently discovered alternative proteome. Curr. Opin. Struct. Biol. 32, 74–80.

Lu, T.C., Leu, J.Y., and Lin, W.C. (2017). A Comprehensive Analysis of Transcript-Supported De Novo Genes in Saccharomyces sensu stricto Yeasts. Mol Biol Evol.

Lynch, M., and Walsh, B. (1998). Chapter 3: Covariance, Regression, and Correlation. In Genetics and Analysis of Quantitative Traits, (Sinauer Sunderland, MA), pp. 35–50.

McLysaght, A., and Guerzoni, D. (2015). New genes from non-coding sequence: the role of de novo protein-coding genes in eukaryotic evolutionary innovation. Philos Trans R Soc Lond B Biol Sci 370, 20140332.

McLysaght, A., and Hurst, L.D. (2016). Open questions in the study of de novo genes: what, how and why. Nat Rev Genet 17, 567–578.

Mouilleron, H., Delcourt, V., and Roucou, X. (2016). Death of a dogma: eukaryotic mRNAs can code for more than one protein. Nucleic Acids Res 44, 14–23.

Neme, R., and Tautz, D. (2013). Phylogenetic patterns of emergence of new genes support a model of frequent de novo evolution. Bmc Genomics 14, 13.

Neme, R., and Tautz, D. (2016). Fast turnover of genome transcription across evolutionary time exposes entire non-coding DNA to de novo gene emergence. Elife 5, e09977.

Neme, R., Amador, C., Yildirim, B., McConnell, E., and Tautz, D. (2017). Random sequences are an abundant source of bioactive RNAs or peptides. Nat Ecol Evol 1, 0217.

Neuhaus, K., Landstorfer, R., Fellner, L., Simon, S., Schafferhans, A., Goldberg, T., Marx, H., Ozoline, O.N., Rost, B., Kuster, B., et al. (2016). Translatomics combined with transcriptomics and proteomics reveals novel functional, recently evolved orphan genes in Escherichia coli O157:H7 (EHEC). BMC Genomics 17, 133.

Ohta, T. (1992). THE NEARLY NEUTRAL THEORY OF MOLECULAR EVOLUTION. Annu. Rev. Ecol. Syst 23, 263–286.

Price, G.R. (1970). Selection and covariance. Nature 227, 520–521.

Reinhardt, J.A., Wanjiru, B.M., Brant, A.T., Saelao, P., Begun, D.J., and Jones, C.D. (2013). De novo ORFs in Drosophila are important to organismal fitness and evolved rapidly from previously non-coding sequences. PLoS Genet 9, e1003860.

Robertson, A. (1966). A mathematical model of the culling process in dairy cattle. Anim Prod 8, 108.

Ruiz-Orera, J., Messeguer, X., Subirana, J.A., and Alba, M.M. (2014). Long non-coding RNAs as a source of new peptides. Elife 3, 24.

Ruiz-Orera, J., Verdaguer-Grau, P., Villanueva-Cañas, J.L., Messeguer, X., and Albà, M.M. (2018). Translation of neutrally evolving peptides provides a basis for de novo gene evolution. Nat. Ecol. Evol. 1.

Schlotterer, C. (2015). Genes from scratch – the evolutionary fate of de novo genes. Trends Genet 31, 215–219.

Soucy, S.M., Huang, J., and Gogarten, J.P. (2015). Horizontal gene transfer: building the web of life. Nat Rev Genet 16, 472–482.

Vakirlis, N.N., Hebert, A.S., Opulente, D.A., Achaz, G., Hittinger, C.T., Fischer, G., Coon, J.J., and Lafontaine, I. (2017). A molecular portrait of de novo genes in yeasts. Mol Biol Evol.

Vanderperre, B., Lucier, J.F., Bissonnette, C., Motard, J., Tremblay, G., Vanderperre, S., Wisztorski, M., Salzet, M., Boisvert, F.M., and Roucou, X. (2013). Direct detection of alternative open reading frames translation products in human significantly expands the proteome. PLoS One 8, e70698.

Wilson, B.A., and Masel, J. (2011). Putatively noncoding transcripts show extensive association with ribosomes. Genome Biol Evol 3, 1245–1252.

Wilson, B.A., Foy, S.G., Neme, R., and Masel, J. (2017). Young Genes are Highly Disordered as Predicted by the Preadaptation Hypothesis of De Novo Gene Birth. Nat Ecol Evol 1, 0146.

Yeh, J. (2006). Real analysis: theory of measure and integration (World Scientific Publishing Co Inc).

Yona, A.H., Alm, E.J., and Gore, J. (2017). Random Sequences Rapidly Evolve Into De Novo Promoters. BioRxiv 111880.

## REFERENCES CITED

Bohrnstedt, G.W., and Goldberger, A.S. (1969). On the Exact Covariance of Products of Random Variables. J. Am. Stat. Assoc 64, 1439–1442.

